# Quantification of Human DNA from Century-Old Archived FFPE Samples for Retrospective Genomic Studies

**DOI:** 10.1101/2025.10.17.681771

**Authors:** V. Zvenigorosky, A. Gonzalez, L. Broisin, J.-L. Fausser, N. Jeanjean, A. Hluszko Pontet, C. Cannet, C. Lamy, C. Keyser, C. Bonah

**Affiliations:** BABEL Laboratory, UMR 8045, Paris, France; Strasbourg Institute of Legal Medicine, Strasbourg, France; Anatomy Unit, Faculty of Medicine, University of Geneva, Switzerland; SAGE Laboratory, UMR7363, Strasbourg, France; Department of Psychiatry, University Hospitals of Geneva, Geneva, Switzerland

## Abstract

**Background:** Formalin-fixed, paraffin-embedded (FFPE) tissue archives are an invaluable resource for genomic research, offering the potential to link genomic data to long-term clinical outcomes. Their utility has however been limited by the degradation caused by fixation and long-term storage, particularly for samples archived for many decades.

**Methods:** This study evaluates a cohort of 79 FFPE tissue blocks collected and archived in 1973 from the Strasbourg Pathological Tissue Archive (SPTA), and a cohort of 51 FFPE tissue blocks from the Geneva Brain Bank (GBB), collected between 1928 and 1971. DNA was quantified using a forensic-grade quantitative PCR (qPCR) assay (QIAGEN Investigator Quantiplex Pro Kit on a Rotor-Gene Q) to precisely measure human DNA concentration, assess degradation, and detect PCR inhibition.

**Results:** A high proportion of the samples yielded human DNA fragments of 80bp and 95bp and very few fragments of 205bp. All organ samples treated with Bouin liquid (n=12) gave poor results, but among samples fixed in formalin (n=117), 58.1% showed over 0.25ng/µL of 80bp human DNA fragments and 37.6% over 1ng/µL. We propose a model to describe the decay of these samples and estimate the proportion of samples that should yield at least a 0.25ng/µL concentration of fragments over 100bp to 45.1%.

**Conclusions:** This work demonstrates that automated extraction methods optimized for FFPE allow for the recovery of usable material even in century-old archived samples with inconsistent conditions of conservation. Our data suggests that the time spent in storage is much less influential on DNA quality than initial fixation time. Crucially, given the fragmented nature of the material recovered (an expected result), future analyses of this material will have to be conducted using next-generation sequencing (NGS) technologies and approaches that rely on short fragments.

## Introduction

Formalin-fixed, paraffin-embedded (FFPE) tissue archives could be one of the most valuable resources in biomedical research (Kokkat et al., 2013; Gaffney et al., 2018; Reingruber et al., 2025), provided they are well documented and sufficiently preserved. For over a century, FFPE has been the standard in tissue morphology analysis (Blum, 1896), leading to the global accumulation of millions of tissue blocks. Where these archives are linked to detailed pathological reports and extensive clinical follow-up data, they constitute an invaluable resource for biomarker discovery or analysis (Grillo et al., 2017), understanding disease evolution (Jiang et al., 2025), and potentially conducting large-scale retrospective studies. The ability to analyze these samples with modern genomic tools would allow researchers to connect the molecular landscapes of the past with long-term clinical outcomes.

Despite this promise, the use of archival FFPE tissues in genomics has been hampered by the deleterious effects on DNA of formalin fixation (among others) (Macabeo-Ong et al., 2002; Steiert et al., 2023) and long-term storage (Von Ahlfen et al., 2007; Guyard et al., 2017). Formalin induces extensive protein-DNA cross-linking (Steiert et al., 2023), complicating extraction. Over time, this can also lead to severe DNA fragmentation (Steiert et al., 2023; Vitošević et al., 2023), in both human and bacterial DNA (Flores Bueso et al., 2020). Furthermore, chemical modifications to DNA bases can introduce sequence artifacts during amplification (Steiert et al., 2023). These challenges collectively result in low yields of poor-quality, fragmented DNA. The limited availability of archived samples and the limited results obtained from degraded FFPE samples have consequently not been conducive to sustained scientific interest. The DNA yields demanded by genomic analyses like whole-genome or whole-exome sequencing are significant (McDonough et al., 2019; Bunduc et al., 2023) and in small collections, attempting these analyses could be a wasted effort.

The SPTA (Strasbourg Pathological Tissue Archive), however, is an archive of several tens of thousands of autopsy cases and hundreds of thousands of biopsy cases, collected between 1945 and 2005 (previous samples ran from 1878 but were discarded before 1945). Each case is associated with sampling records, histological reports, FFPE (Formalin-Fixed Paraffin-Embedded) blocks, stained and unstained microscope slides and, in some cases, medical records. Crucially, the availability of DNA in some SPTA samples has already been demonstrated (Zvenigorosky et al., 2024). The second archive presented here, the GBB (Geneva Brain Bank), is an actively expanding collection of 10,000 formalin-fixed brains and their associated 100,000 FFPE blocks (with a smaller number of other organs) and 200,000 microscope slides (Kövari et al., 2011). Each case of the GBB is accompanied by neuropathological diagnostic reports and clinical records that allow us the same detailed comparisons as in the SPTA. The collection was started in 1901 and is curated at the Belle-Idée Psychiatric Hospital of the University Hospitals of Geneva. For these two archives, this wealth of documentation and material therefore covers historical data, archival data but also unequaled analytical potential for molecular biology.

The present work aims at going further than the punctual analysis of archived FFPE samples, and at estimating the proportion of viable samples in these archives, in order to propose guidelines for other collections, and evaluate the potential of the archive more precisely. We selected 79 FFPE samples from 1973, the beginning of block standardisation in Strasbourg, as a benchmark after which it can be surmised that samples will have similar or better DNA yields, given the technical improvements and the significant increase in homogeneity, both material and methodological. These improvements are namely the exclusive use of Formalin and not Bouin’s fluid, the end of beeswax use, control over fixation time and long-term storage in custom-made shelving. From the GBB, where methods are more homogenous, and the collection less extensive, we selected 51 FFPE samples within a cohort of syphilis patients on which bacterial DNA analyses are planned, as part of the ArchiMED project (SNSF 219457, ANR-23-CE45-0028). Selected cases were collected between the late 1920s and 1970s, a period of intensive sample collection that overlaps with numerous neurosyphilis cases.

The present work shows that the proportions of FFPE blocks yielding usable DNA in sufficient quantities allows for the planning and execution of retrospective genetic studies on the SPTA and the GBB, and likely other archives, without the necessity of performing extractions on a prohibitively large number of samples. Results are also consistent with the conclusion that fixation time is the critical factor in determining DNA availability, rather than long-term storage duration.

## Materials and Methods

### Ethical Statement

This study was approved for the SPTA by the Strasbourg Faculty of Medicine Ethics Committee under number CE-2025-10, as part of the ArchiMed project (ANR-23-CE45-0028). All work adheres to strict anonymization processes, which are facilitated by the use of samples that had been previously coded as part of the standard medical analysis process.

The GBB is registered with the Cantonal Research Ethics Commission (CCER)/Swissethics under protocol number 2017-01937. The present study is part of the ArchiMed project (Swiss National Science Foundation project no. 219457), approved by the CCER under reference number 2024-01790, with the same constraint on anonymization, which is facilitated by the original coding.

### Sample Cohort

We selected 79 FFPE tissue blocks from different cases in the SPTA (Supplementary Table S1), all collected and archived in the year 1973. Our samples were selected from large drawers containing smaller boxes of six to twelve blocks of similar sizes. From each box, we selected one case and collected all the blocks corresponding to that one case. We only subjected one block per case (with a few exceptions) to sampling for further analyses and we did not repeat analyses, in an effort to provide a “first instance” approach to the availability of DNA. Given histopathological reports, the identification of organs and pathologies is very straightforward and easily standardized in the SPTA. Detailed statistics are presented in the Results section.

From the GBB, 26 patients diagnosed with neurosyphilis were selected based on confirmed pathological diagnosis, documented clinical symptoms, historical time period, and the preservation quality of the available material. Patients from each decade between 1928 to 1971 were included. For each case, the corresponding formalin-fixed whole brains were available for potential further analysis. A total of 51 FFPE blocks were retrieved from the 26 selected patients, primarily from the central nervous system (various brain regions and the spinal cord), along with a limited number of samples from other organs, fixed mainly with Bouin’s solution. The decision not to include a larger number of Bouin-fixed samples (and none from the SPTA) is motivated by the poor results expected. Preliminary tests in the SPTA using disparate methods and FFPE in poor storage conditions with unclear technical characteristics have returned very low concentrations of human DNA, and highlighted the necessity of focusing first on this later period of FFPE collection, to control for known parameters. Bouin fixed blocks are in any case the small minority of either archive. Since in the early decades of the collection, the anatomical origins of sampled tissues were not labeled on the blocks (but on inserts placed inside the box), we consulted the corresponding archived histological slides to identify the organ or regions collected in the selected blocks. Specific brain regions were identified either by direct anatomical labeling on the slides or, using the Von Economo – Koskinas nomenclature (Triarhou, 2007; García-Cabezas et al., 2020) often used and written on the slides. Archival slides were also compared with newly prepared hematoxylin and eosin (H&E) stained sections obtained from the same blocks to confirm the identification.

### Collection, fixation and fixation time

The SPTA is the deposit of FFPE samples that was produced during the routine work of the Strasbourg Institute of Pathology during the last 150 years (although most of the surviving material was produced after 1945). All FFPE blocks were used to produce microscope slides intended for anatomopathological analysis, either following a medical autopsy or in a clinical context. Early samples were often treated with Bouin’s fluid, but formalin and then buffered formalin use was becoming the standard in Strasbourg in 1973. While autopsy FFPE samples fixation times vary greatly, from hours to years, the blocks presented here were collected from the clinical section and were produced under time constraints, since they were used to inform the treatment of living patients. Consequently, the samples presented here were immersed in formalin for less than seven days, with many under one day. While fixation time is not indicated in the reports, we have based our analysis on the time between sampling and the histological report. However, our knowledge of historical methods, as witnessed and practiced by one of the co-authors of the present work, would suggest that very few samples have spent several days in formalin, given the relative automation of the fixation process by 1973.

The GBB is a brain biobank containing FFPE samples obtained from medical autopsies from the Belle-Idée hospital since 1901. Autopsies were performed in the neuropathology unit of the Psychiatric hospital, and since 1971 in the geriatric hospital as well. Tissue samples comprising formalin-fixed tissues, FFPE blocks and associated histological slides, have been prepared since 1901. The frequency of autopsies in the GBB was lower than the frequency of biopsy in the SPTA (these covering any and all cases). Fixation times for GBB samples are longer (and increase with time) than those for SPTA samples, as SPTA autopsies are performed for diagnostic purposes and therefore require faster processing, whereas GBB autopsies are less time-sensitive. This gave the collectors of the GBB more leeway to leave brains (or other organs) in formalin for extended periods. The fixative used was typically 15% unbuffered formalin (equivalent to ∼5% formaldehyde), for 3 to 4 weeks. Brains were then transferred in 5% formalin for long-term storage. The level was regularly monitored to assess evaporation and specimen condition. Today, the formalin is renewed approximately every two years to maintain optimal preservation conditions.

A limited number of organs (including liver, lungs, optic nerve, pituitary gland, rectum, thyroid, pancreas, and aorta) were also sampled in the collection, although not on a systematic basis, until after the second half of the 20th century. These specimens were typically retained when considered of pathological relevance and were fixed with Bouin’s fluid.

### Paraffin embedding and labelling

Formalin-fixed samples, both in the SPTA and GBB were dehydrated through graded ethanol (or possibly in some cases methanol in the SPTA) series. Clearing was performed using toluene, benzene or xylene. For the GBB, clearing was originally performed using toluene or benzene. These solvents were replaced by xylene around 2010, because of their limited commercial availability and higher toxicity. For the SPTA, the detailed history of practice is not documented but ethanol and xylene have progressively become the standard.

In the GBB, the embedding medium initially used for tissue inclusion consisted of a 1:1 mixture of two types of paraffin, one of which was supplemented with beeswax. This paraffin was manually filtered prior to use to remove impurities. Before 1970 in the SPTA, a 2:1 mix of paraffin and beeswax was used. The samples presented here, from 1973, were embedded in paraffin alone, marking the the end of beeswax use and the beginning of the practices standardization. In Geneva, the introduction of synthetic paraffin occurred later, in the early 2000s. GBB case numbers have consistently been indicated either by paper label affixed to the container or by an insert placed directly within the sleeve throughout the period, while 1973 SPTA blocks had the case number engraved by the technician on one or two sides. In the GBB, the patient number also began to be engraved on the blocks in the 1990s.

### Sample storage *in situ*

SPTA clinical samples (around 800,000 cases, likely over a million blocks) are stored in a large basement room of the Strasbourg Institute of Pathology, which undergoes limited temperature variations but where humidity is often high. The samples are placed on custom-made wooden and metal shelves that host three types of material: (1) the older blocks of diverse shapes and sizes, in small cardboard boxes (10-20 blocks), organized in larger cardboard boxes (of a few dozen small boxes). These FFPE samples were moved into their current configuration in the early 1960s. (2) Partially size- and material-standardized blocks are organized in metal shelves where each drawer contains small open-top cardboard boxes where the FFPE samples are organized and sorted by case number. Each small box contains 10-15 blocks and each drawer contains a few dozen boxes. The samples presented here are stored in these metal shelves. (3) Standardized cassette FFPE samples are stored mainly in wooden shelves built on the model of the previous metal shelves.

Concerning the GBB, paraffin-fixed tissues and fixed brains were initially stored in the attic of the laboratory building of Belle-idée Psychiatric Hospital (Geneva). The earliest preserved FFPE tissues and fixed brains in the collection date back to 1926, as earlier specimens were discarded in 1925. Infrared lamps were installed to prevent temperature fluctuations and cooling of the attic and the area was ventilated during summer months. However, some of the oldest FFPE blocks partially melted. In the 1980s, the tissues were relocated to the basement level of the same building due to increasing concerns about the structural load capacity of the upper floors, given the cumulative weight of the growing collection. Since that time, the collection has remained in the basement. The tissues are stored at room temperature under controlled conditions, in ventilated rooms.

The FFPE specimens were initially stored in matchboxes. In the 1950s, these were progressively replaced by microscope slide and coverslip boxes, and from the 2000s onwards, sealed plastic sleeves became the standard storage method. Each patient has between 1 and 20 blocks on average.

Over the years, the shelving systems have been progressively replaced: the earliest cases are stored on wooden shelves in the first room, while more recent cases are housed on electro-galvanized metal shelving units in the three adjoining rooms.

### Tissue Preparation for the present study

For each of the 79 FFPE blocks from the SPTA, 10mg to 20mg of tissue were carved out using a single-use scalpel. Block surfaces were first decontaminated using ultrapure water, and the upper layers of paraffin removed. The scalpel was then decontaminated using DNA Away (Thermo Fisher Scientific) and cleaned with ultrapure water. Depending on the tissue, certain areas were preferred, under the supervision of a histologist (the pertinence of these decisions is discussed in the last sections of this work). The amount of tissue collected (in mg) was recorded.

Each of the 51 FFPE blocks from the GBB was first trimmed to remove upper sections, and serial sections of 24μm thickness were then obtained using a RNase and DNase free microtome. For each patient, sections were placed into 1.5 mL sterile tubes for DNA extraction, under DNase-free conditions. Depending on the block size, one to three sections were placed in each tube. To measure the initial tissue volume, an image of each block was taken alongside a calibrated scale. The surface area (in mm^2^) was measured using Fiji (ImageJ), and the tissue volume (in mm3) was calculated by multiplying the measured area by the section thickness. The resulting recorded volumes ranged from 0.35 mm^3^ for the smallest sample to 13.96 mm^3^ for the largest.

### DNA extraction

DNA extraction for the SPTA was performed on the QIAGEN EZ2 Connect automated platform, according to the manufacturer’s protocols, using the EZ2 AllPrep DNA/RNA FFPE Kit (QIAGEN).

DNA extraction for the GBB was performed on the Promega Maxwell® CSC 16-channel instrument (operated in Research mode), using the XtractAll FFPE DNA/RNA Kit or Xcelerate DNA FFPE Kit. All steps were carried out according to the manufacturer’s protocols, except for the deparaffinization step, where xylene treatment was used instead of mineral oil. Samples were incubated twice with xylene for 5 minutes at 56°C under agitation (400rpm), followed by maximum speed centrifugation. The tissues were then washed in three successive ethanol baths, each followed by centrifugation and finally air-dried at 56 °C for 5 minutes or until completely dry, without agitation.

### DNA Quantification and Quality Control

Considering the known contamination of the material, quantification had to target human DNA specifically, rather than the overall content of the FFPE samples (McDonough et al., 2019). Quantification was performed using the QIAGEN Investigator Quantiplex Pro Kit on a Rotor-Gene Q real-time PCR cycler, according to the manufacturer’s protocols. Three separate experiments, each including two blank controls, covered all 129 samples.

### Contamination

Qiagen’s dedicated spreadsheet tool proposes an estimation of “mixture”, which is based on the discrepancy between the “Human DNA” quantity measured and the “Male DNA” quantity. Since these quantities are based on fragments of 80bp and 95bp respectively, it is surmised that for a male individual, they should be roughly equivalent (both fragments are short). In the context of our study however, this method proved unusable, due to an average fragment size around these values. In some cases, males showed satisfactory autosomal DNA quantification but no Y-chromosome DNA quantification. We therefore estimated DNA on a case by case basis, relying on the known sex of individuals and the likelihood of contamination during sampling or preparation given different organs and the specific conditions of their handling.

The historical collection process in the SPTA presented four distinct risks of contamination, in order of decreasing likelihood: (1) the organs were cut on a single board that was cleaned once a day, after dozens or hundreds of samples had been cut; (2) the organs were handled by technicians who did not wear protective equipment beyond gloves; (3) best practices were not strictly followed during surgery, especially in the earlier periods and (4) the automated fixation apparatus in Formalin was designed to handle several samples at once. Some of these circumstances would lead to cross-contamination between samples (although cross-contamination in a formalin bath is unlikely) and the second instance would lead to contamination by the technician or technicians. We have endeavored to test for these events in the data presented here, although experience indicates that only the risk of cross-contamination is likely to be significant.

The nature of the GBB makes it much less susceptible to sample cross-contamination. Brain collection during autopsy was not a daily event and more time and care was invested in each case. Each brain was fixed in isolation in a bucket of formalin and all have remained isolated.

### Statistical Analysis

Descriptive statistics and modelling were performed with R 4.1.2 and custom python 3.12 scripts (using modules numpy, matplotlib, scipy, scikit, seaborn and pandas).

## Results

### Overall DNA Quantification Success, Degradation and Inhibition

The GBB samples included 12 Bouin-fixed organ samples. Of those, 7 did not yield any DNA, 3 yielded under 0.1ng/µL of the 80bp fragment, and 2 yielded under 0.2ng/µL. Although this small number of samples does not permit general conclusions, it is compatible with the expectation that Bouin’s fluid causes more extensive degradation to the DNA (De Martino et al., 2022). We have excluded these 12 samples from further statistical analyses but we will include the most successful ones in future molecular analyses. Additionally, 1 formalin-fixed sample from the GBB presented incoherent values for the 80bp fragment (over 200ng/µL) and was excluded from the analyses.

Out of the 117 FFPE samples retained for statistical analyses (Figure 1), when considering first the 80bp fragment (the shortest), 19 (16.2%) yielded no DNA or less than 0.005ng/µL, 30 (25.6%) yielded between 0.005ng/µL and 0.25ng/µL, 24 (20.5%) yielded more than 0.25ng/µL and 44 (37.6%) yielded more than 1ng/µL. These results are presented in Figure 1 (section A).

**Figure 1:**
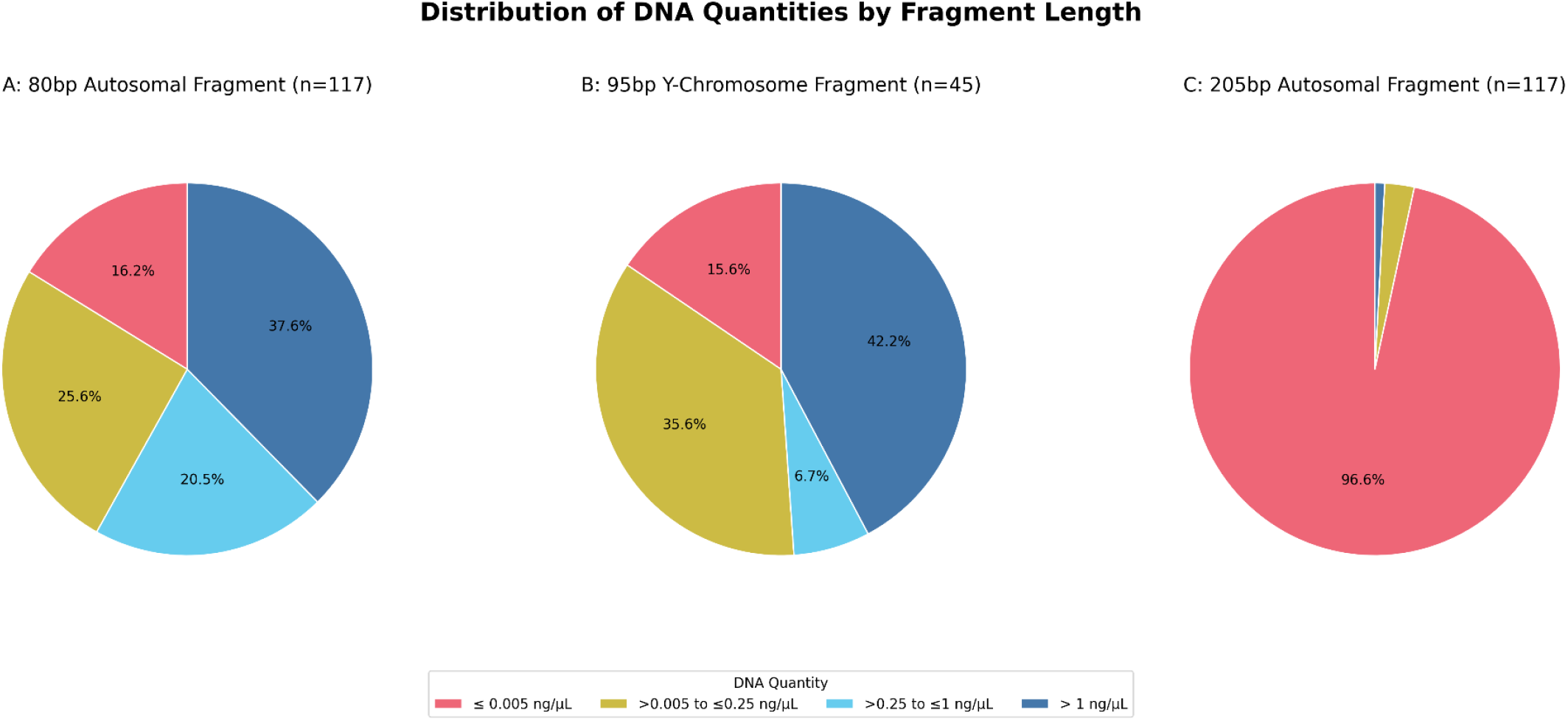
Distribution of DNA Quantities by Fragment Length

For the 45 FFPE samples from males, we computed the quantities for the 95bp fragment. 7 (15.6%) yielded no DNA or less than 0.005ng/µL, 16 (35.6%) yielded between 0.005ng/µL and 0.25ng/µL, 3 (6.7%) yielded more than 0.25ng/µL and 19 (42.2%) yielded more than 1ng/µL. These results are presented in Figure 1 (section B).

Our analysis includes the amplification of the 205bp fragment for all 117 samples. For this last fragment, 113 (96.6%) yielded no DNA or less than 0.005ng/µL, 3 (2.6%) yielded between 0.005ng/µL and 0.25ng/µL and 1 (0.9%) yielded more than 1ng/µL. These results are presented in Figure 1 (section C). This implies that the degradation index, which is a ratio of the quantity of 80bp fragments and the quantity of 205bp fragments, could only be calculated for 4 samples. No inhibition flags were raised by the software.

### Breakdown by archive

A Mann-Whitney U test was performed to determine if there was a statistically significant difference in the quantity of the 80bp autosomal fragment between formalin-fixed samples from the GBB and SPTA archives. The results of the test indicate that there is no statistically significant difference in the DNA quantities between the two groups (U = 1448.5, p = 0.76). The high p-value suggests that any observed differences in the median DNA yields are likely due to random sampling variation rather than a true, systematic difference in preservation quality between the two archives.

This statistical conclusion is supported by the visual representation in the comparative box plot. The plot shows that the distributions of DNA quantities for both the GBB and SPTA archives largely overlap, indicating similar overall performance. While the median quantity for the SPTA samples is slightly higher, the interquartile ranges are comparable, and the GBB archive contains individual samples with higher maximum yields. Taken together, the statistical test and the visualization confirm that for formalin-fixed tissues, both archives provide a comparable source of amplifiable DNA.

### Breakdown for each organ category

To assess the influence of tissue type on DNA preservation, we compared the quantities of the 80bp autosomal fragment across different organs. A Kruskal-Wallis H test performed on the seven most represented organ groups (Figure 2) revealed a statistically significant difference in DNA yield among them (H = 25.29, p = 0.0003), indicating that the organ of origin is a critical factor in the recovery of amplifiable DNA.

**Figure 2:**
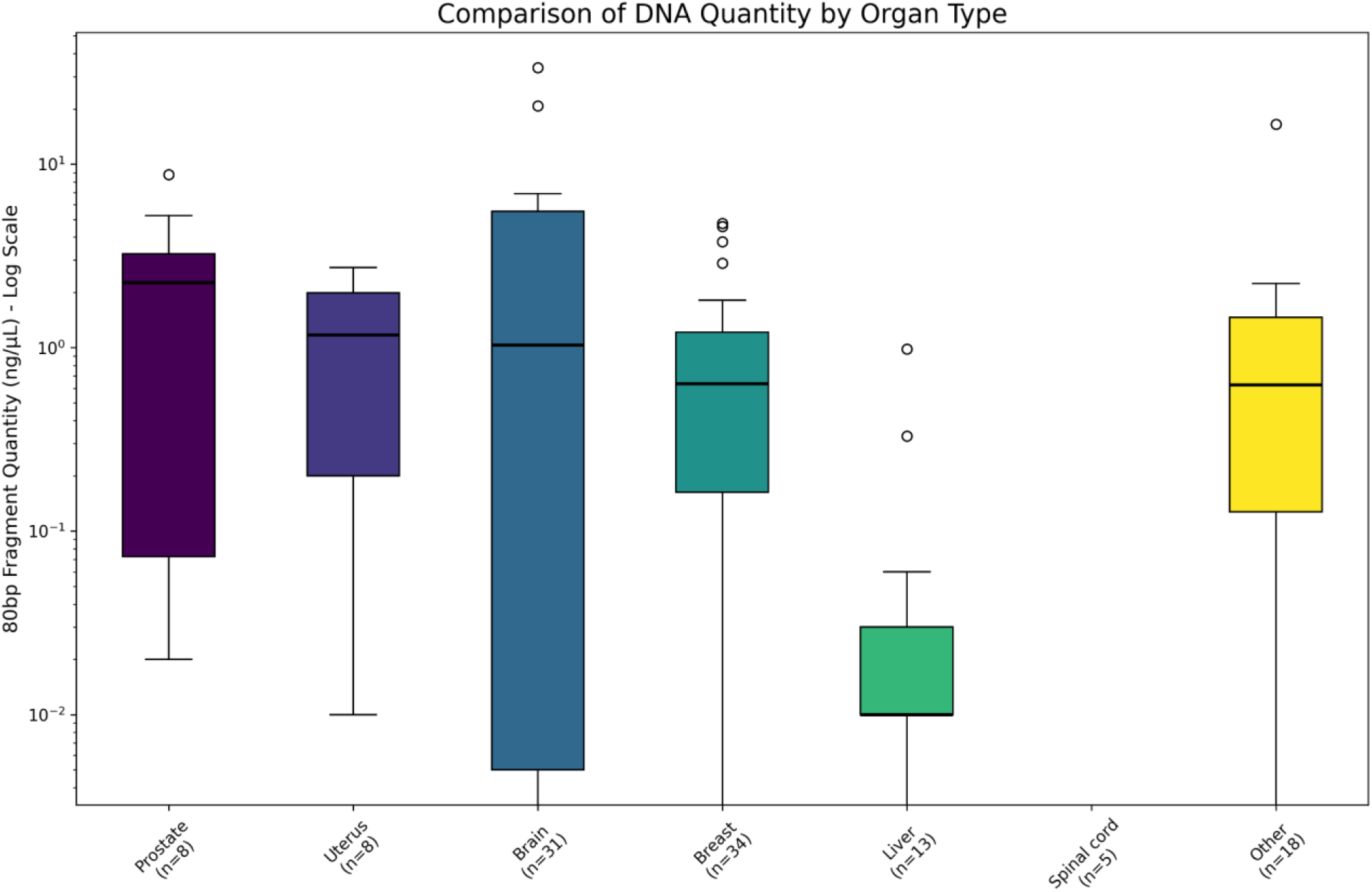
Comparison of DNA Quantity by Organ Type

A post-hoc Dunn’s test with Holm-Bonferroni correction was conducted to identify the specific sources of this variation. The pairwise comparisons revealed two distinct clusters of tissues based on preservation quality. The first cluster, consisting of Liver and Spinal cord, showed consistently poor DNA yields with no significant difference between them (p = 1.00). The second, larger cluster, which includes Brain, Breast, Prostate, Uterus, and the “Other” category, demonstrated significantly better preservation, with no statistical differences in DNA yield among any of the tissues within this group (p ≈ 1.00 for all intra-group comparisons). The key finding is the highly significant difference between these two clusters, where tissues in the well-preserved group consistently yielded more amplifiable DNA than those in the poorly preserved group (p < 0.05 for most inter-group comparisons). This confirms that tissue composition is a primary determinant of DNA stability in FFPE blocks.

### Effects of fixation and fixation time

To investigate the impact of fixation duration on DNA yield, the GBB and SPTA archives were analyzed separately due to their fundamentally different fixation protocols. For the SPTA cohort, where fixation time represents a recorded maximum between sampling and analysis, a chi-squared test revealed no statistically significant association between the duration of fixation and the distribution of DNA yields (χ^2^ = 9.64, p = 0.65). As expected, successful and unsuccessful outcomes were observed across all fixation time categories. This suggests that within the short timeframes typical of the SPTA protocol (0-10 days), the precise duration of fixation is not a primary determinant of DNA preservation, especially since tissue sample size and nature directly affects the penetration of formalin.

In contrast, the GBB archive, characterized by a much longer and more variable minimum fixation period related to autopsy procedures, showed a highly significant association between fixation time and DNA yield (χ^2^ = 17.84, p = 0.0013). The stratified analysis (Figure 3) clearly demonstrates this trend: samples fixed for less than one month show a high proportion of successful DNA recovery, whereas yields drop sharply for samples fixed between one month and one year. Critically, no samples fixed for longer than one year yielded usable DNA (Figure 3 excludes one sample fixed for more than 34 years). This strong negative correlation underscores that prolonged immersion in formalin, as practiced in the GBB protocol, is a major cause of DNA degradation, severely limiting the viability of samples subjected to long-term fixation.

**Figure 3:**
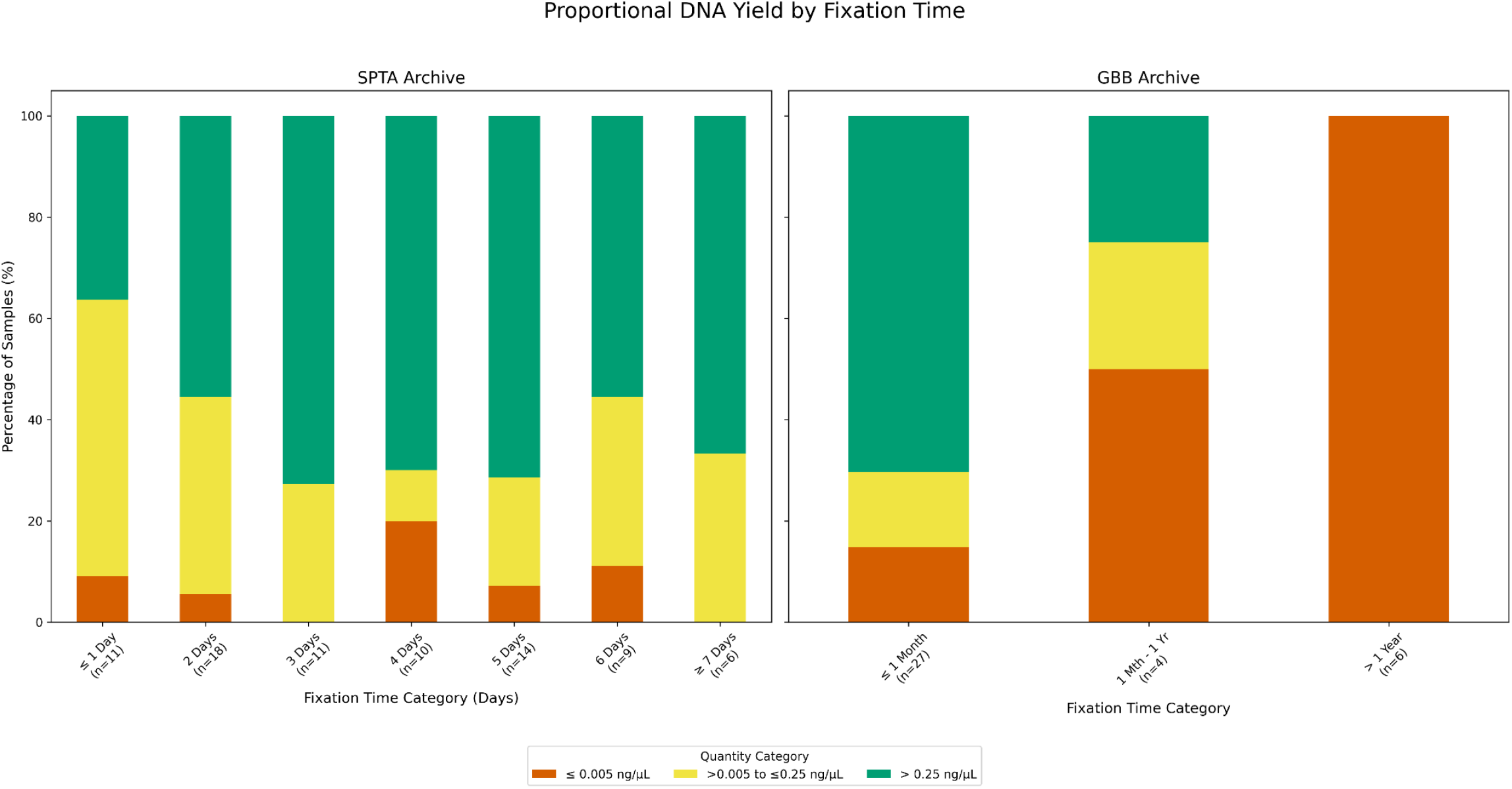
Proportional DNA Yield by Fixation Time

### Correlations with the mass or volume of tissue sampled

To determine if the amount of starting material influenced DNA recovery, we analyzed the relationship between the initial quantity of tissue processed and the final DNA yield for each archive separately. For the SPTA cohort, where tissue was measured by mass, a chi-squared test revealed no statistically significant association between the amount of tissue sampled and the distribution of DNA yields (χ^2^ = 4.81, p = 0.31). This indicates that for the SPTA samples, using more or less tissue within the tested range did not have a predictable effect on the success of the DNA quantification (Figure 4). In contrast, the analysis of the GBB cohort, where tissue was measured by volume, showed a statistically significant association between the starting volume and the final DNA yield (χ^2^ = 10.51, p = 0.03). Notably, this relationship was negative; samples with a larger starting volume resulted in poorer DNA yields. Since the qPCR analysis did not indicate PCR inhibition for these samples, this finding may suggest an upstream issue, such as an inhibition or saturation effect during the automated DNA extraction process itself when processing larger tissue volumes. This result implies that for GBB samples, increasing the amount of starting tissue did not improve, and may have even hindered, the efficiency of DNA recovery.

**Figure 4:**
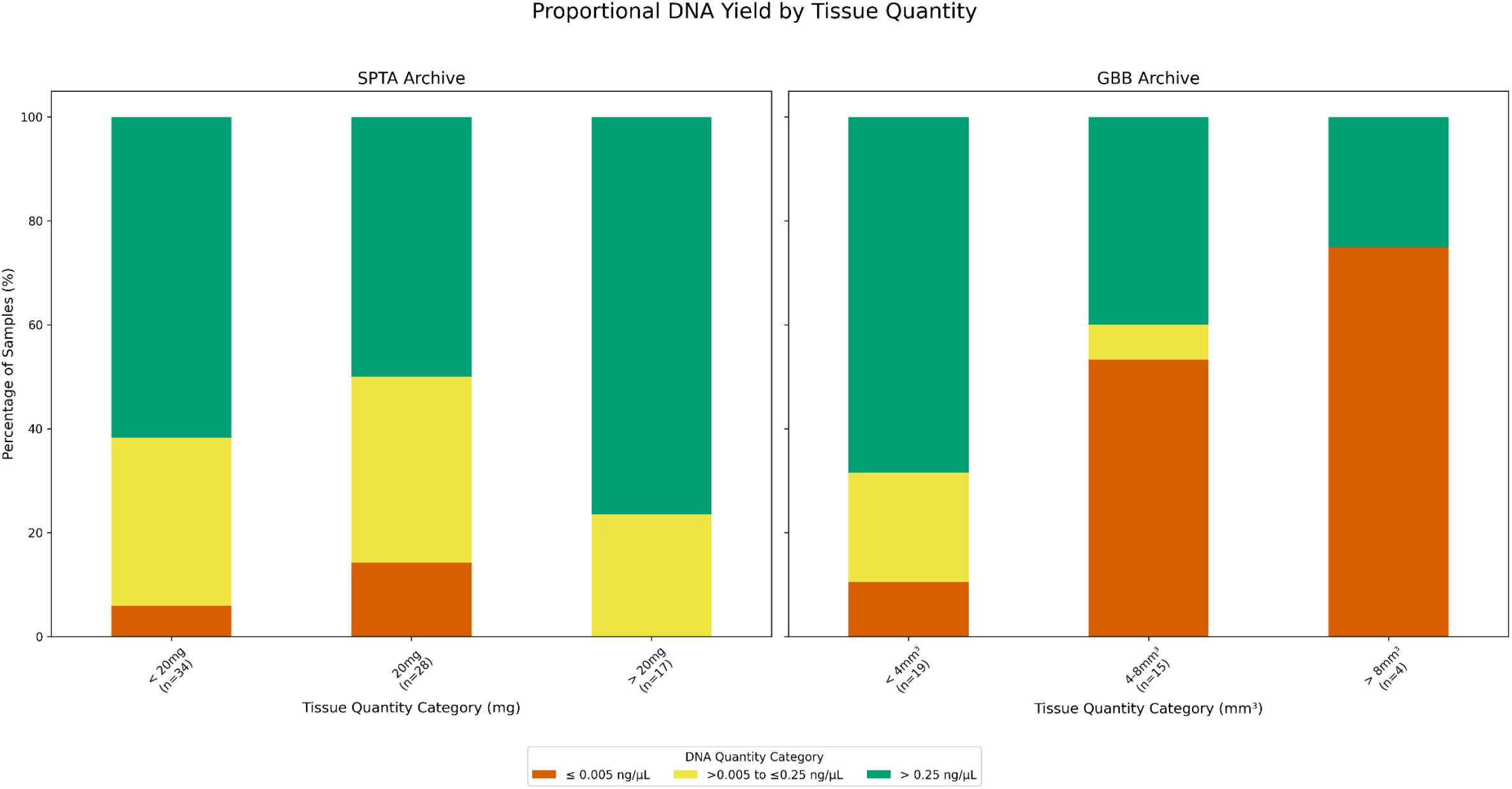
Proportional DNA Yield by Tissue Quantity

### Contamination and errors

Controls showed no signs of contamination, but one previously mentioned sample presented the improbable value of over 200ng/µL for the short fragment. This sample corresponded to a female patient and was unremarkable regarding fixation time (21 days, like 64% of the GBB), organ (the left occipital pole of the brain, similar to most of the GBB), fixator (Formalin) or year of sampling (1949 within our range of 1928 to 1971). The volume of tissue sampled was among the highest for the GBB, but not significantly so (10.55mm^3^ with the average was 5.52mm^3^ and the range 0.35mm^3^-13.96mm^3^). The quantity of 95bp fragments amplified was <0.005ng/µL (consistent with a female) and the quantity of 205bp fragments amplified was 1.77ng/µL. Given the lack of any indication of error, but considering the unlikelihood of such a high quantity of contaminant autosomal DNA in the sample (Daly et al., 2012), we took the decision to exclude that sample until other such events can be identified and characterized.

Given the degradation of the material, the only definitive detection of contamination is the case of female samples presenting male DNA. Only three female samples (all breast tissue from the SPTA) showed such contamination. Comparing the quantity of the 95bp Y-chromosome fragment to the autosomal 80bp fragment in all males and the three contaminated female samples (Figure 5) shows that two of the latter could not have been distinguished from males if the information was previously unknown. One female sample showed a very small quantity (0.01ng/µL) of the 95bp fragment (ratio ∼143) and could therefore not be mistaken for a male sample. It should be noted that the four samples of unknown sex (three placentas and one embryo) showed no male DNA, but the quantities obtained for the 80bp fragment being low for those samples (0, 0.01, 0.11 and 0.46ng/µL), we could not ascertain whether the samples were male or female.

**Figure 5:**
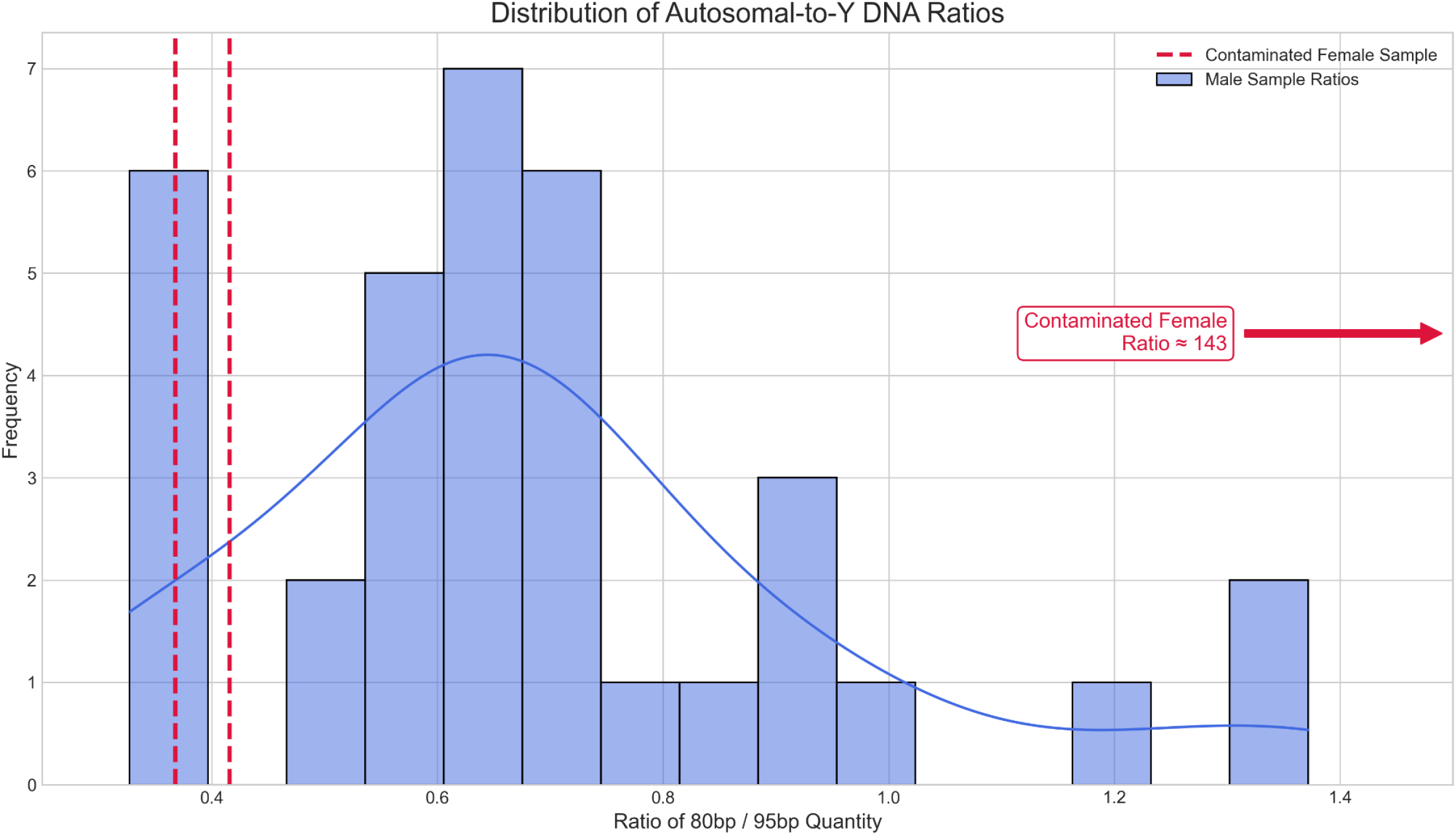
Distribution of Autosomal-to-Y DNA Ratios

## Discussion

### Degradation models

We modeled the probability of successful amplification as a function of fragment length. Using a simple exponential decay model, as seen in ancient DNA studies (Allentoft et al., 2012), resulted in a poor fit to our observed data. This finding is significant, as it suggests the degradation pattern in these FFPE archives is not a smooth, continuous process. Instead, the data was accurately described by a weighted logistic decay model (Figure 6), which captures a sharp drop-off in amplification success around a critical failure point.

**Figure 6:**
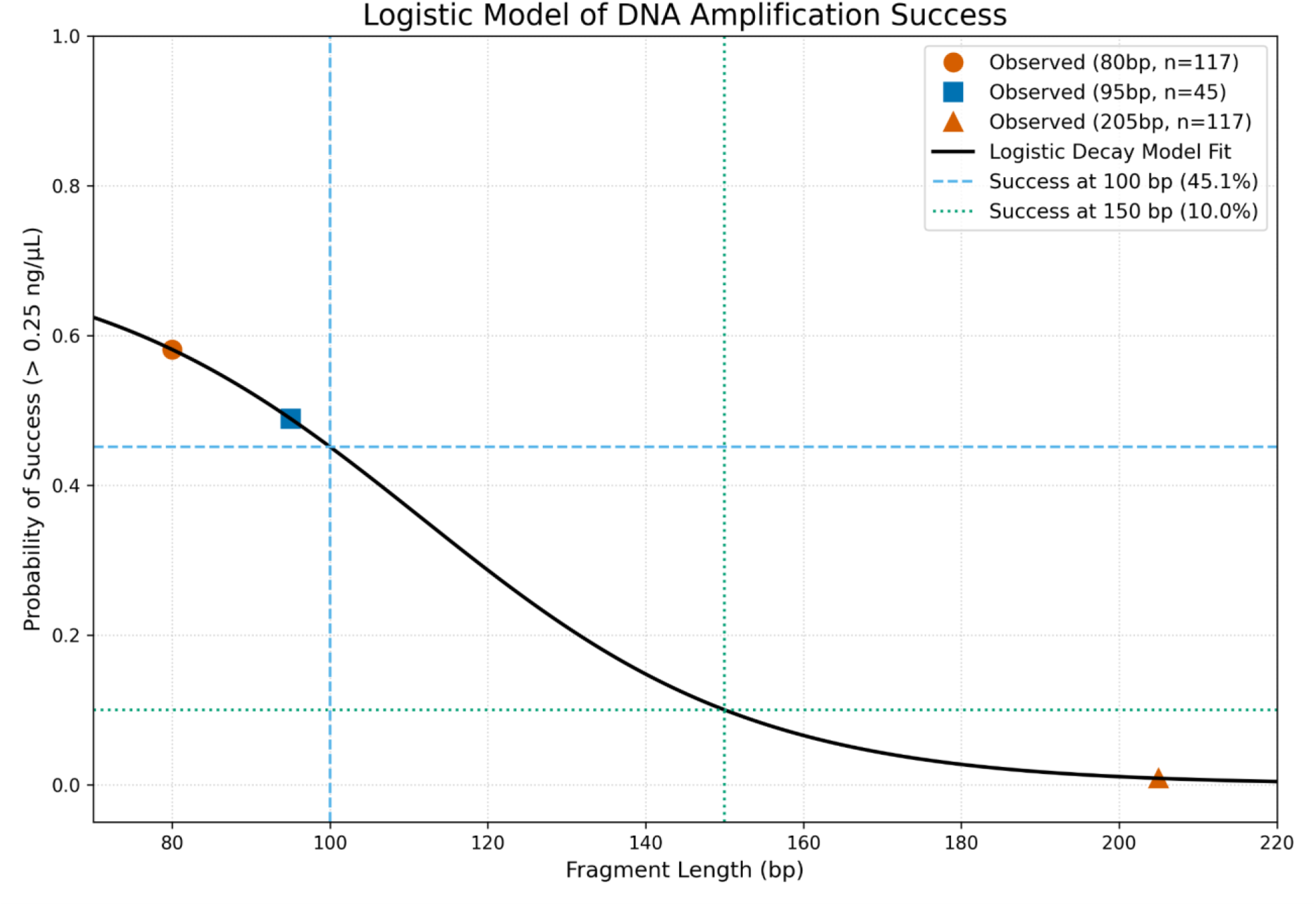
Logistic Model of DNA Amplification Success

We do not propose that this logistic decay pattern is a model of degradation rate over the past 50 years, but rather a descriptive curve fit of the data, showing the consequences of the initial, heterogeneous degradation that occurred during formalin fixation. This situation is expected, as some degradation occurs while samples are being kept in the archive (over the decades), but initial fixation has a much greater effect.

By establishing a robust relationship between fragment length and amplification success, we can estimate the viability of future assays on this or similar archives. Based on our model, a hypothetical assay targeting a 100 bp fragment would have a predicted success rate of 45.1% for yielding >0.25 ng/µL of DNA. For a longer 150 bp fragment, a length common in many NGS panels, the predicted success rate drops to 10.0%.

### Usability in downstream NGS analyses

The primary objective of quantifying DNA from archival FFPE samples is to determine their suitability for downstream next-generation sequencing (NGS) applications. The utility of a sample for NGS is defined by two key metrics: the total mass of amplifiable DNA and the integrity (fragment length) of that DNA. While historically a significant challenge, our results demonstrate that a substantial portion of these century-old FFPE samples meet the requirements of modern, sensitive NGS workflows.

Contemporary NGS library preparation kits, particularly those designed for damaged DNA, often require a total input of 1-10 ng of DNA. Our quantification data shows that a significant fraction of the archive can meet this threshold. When considering a practical cutoff of >0.25 ng/µL, 58.1% of the 117 non-Bouin fixed samples were successful. More stringently, 37.6% of samples yielded over 1 ng/µL from the 80bp autosomal target alone, indicating that many samples contain sufficient material for demanding applications like whole-exome sequencing or complex targeted panels. The success of the 80bp assay is particularly relevant, as its short amplicon length serves as an excellent proxy for the fragment sizes typically targeted during NGS library construction.

Beyond simple quantity, our logistic decay model provides a predictive framework for assessing the archive’s potential for assays targeting different fragment lengths. The model predicts that for an assay requiring a yield of >0.25 ng/µL, approximately 45.1% of samples would be viable for a 100 bp target, and this rate drops to 10.0% for a 150 bp target. This allows for a data-driven approach when selecting samples for specific NGS panels, balancing the desired fragment length against the expected sample success rate. While our analysis identified low-level male DNA in a small number of female samples, the distinct separation of these samples from the male population suggests that such contamination could likely be identified and bioinformatically filtered out during post-sequencing analysis.

It is crucial to note, however, that this functional quantification addresses only the first two hurdles for NGS: DNA quantity and fragmentation. It does not provide information on the chemical quality of the DNA sequences themselves. Formalin fixation is well-documented to cause chemical modifications, primarily cytosine deamination, which can lead to C>T substitutions in the final sequencing data (Einaga et al., 2017). Therefore, while our results provide a robust foundation for sample selection and cost estimation, subsequent pilot sequencing studies will be essential. Such studies are necessary to quantify the rate of formalin-induced artifacts and to validate bioinformatic strategies for filtering these errors, ensuring the ultimate fidelity of any genomic data generated from these invaluable historical archives.

### Organ-specific sampling guidelines

Several factors are crucial for optimizing yield and quality, particularly for sample selection in archives. Our results suggest that the spinal cord, when fixed alongside the brain, is highly susceptible to overexposure to formalin, leading to complete DNA degradation. Similarly, our experience indicates that for breast cancer samples, DNA should be preferentially extracted from the protein/tissue area rather than the lipidic region of the blocks to maximize recovery.

For other tissues, general guidelines apply: for percutaneous biopsies, the entire sample should be utilized (given the small size of samples). In prostate cancer, human DNA can be sampled broadly, though for tumor analysis, multiple samples from various locations may be necessary, with histological guidance. For brains, human DNA can be sampled from any region, or specifically from areas of inflammation in cases of syphilis. Placenta or embryo samples can be taken from any part of the tissue. Bones may present accessibility challenges due to decalcification, though they are uncommon in the SPTA and absent in the GBB.

## Conclusion

This preliminary study of 130 FFPE samples from the Strasbourg Pathological Tissue Archive and the Geneva Brain Bank, with some specimens dating back to 1928, demonstrates the feasibility of conducting retrospective genomic studies on very old and variably preserved collections. Our results show that despite significant DNA fragmentation, a substantial proportion of formalin-fixed samples (58.1%) yield enough short fragment DNA (>0.25 ng/µL at 80bp) to meet the input requirements of certain modern next-generation sequencing workflows.

We identified and confirmed several critical factors that determine DNA recovery success. Organ type proved to be a primary determinant, with tissues like brain and breast showing significantly better preservation than liver or spinal cord. Furthermore, while the short fixation times typical of the SPTA clinical workflow did not significantly impact yield, the prolonged multi-month formalin immersion common in the GBB protocol was found to be more variable in its effects, especially concerning whole brain fixation. We propose a simple degradation model that takes into account initial fixation heterogeneity as the principal cause of DNA damage. The predictions made by this model are now testable in future studies.

These results decisively demonstrate that there is great scientific value in large medical archives of FFPE. Not all samples will yield usable quantities of DNA, but the proportions of available material are sufficient so that the number of necessary attempts is not prohibitive. For some NGS approaches, around 50% of samples would yield sufficient material. We therefore propose that new technological advances have made this kind of material accessible for genetic and genomics research.

## Supplementary Materials

Supplementary Table S1: Samples in the cohorts and quantification results

